# Plant provenance affects pollinator network: implications for ecological restoration

**DOI:** 10.1101/2020.11.26.399493

**Authors:** Anna Bucharova, Christian Lampei, Malte Conrady, Emilia May, Janis Matheja, Michael Meyer, David Ott

## Abstract

The selection of plant provenance for ecological restoration is an intensively debated topic. Throughout this debate, arguments mostly focus on plant performance, but little attention is paid to the effects of provenance on other members of the restored ecosystem. On the other hand, in restoration projects that focus specifically on supporting interacting biota, for example flower stripes among fields to support pollinators, the provenance choice is often not considered, partly because the effect of provenance on pollinators is unknown. In this pioneering case study, we tested whether pollinators differentiate between experimental plant communities of different provenances.

We established experimental plant communities with the same species composition but with plants originating from three different provenances. We then recorded plant phenology and observed pollinators and flower visitors interacting with these experimental communities and related the pollinator visitation to the provenance identity.

The provenances of the experimental plant communities had a strong and significant effect on the diversity and abundance of flower-pollinator interactions, with one provenance interacting twice as often as the other two provenances. The effect was driven by the differences in flowering phenology among provenances.

**Synthesis and applications:** Plant provenances substantially differ in their interactions with local pollinators. Therefore, the selection of plant provenance should be considered when planning restoration projects for the support of pollinators.

## Abstract in German

Die richtige Auswahl der Herkunft von Pflanzen für die ökologische Renaturierung ist ein intensiv diskutiertes Thema. In dieser Debatte liegt der Fokus in erster Linie auf der erfolgreichen Etablierung der Pflanzen, weit weniger Aufmerksamkeit erhält die Auswirkung der Herkunft auf andere Mitglieder des wiederhergestellten Ökosystems. In Renaturierungsprojekten, wiederum, die sich speziell auf die Unterstützung interagierender Biota konzentrieren, z.B. Blühstreifen zwischen Feldern zur Unterstützung von Bestäubern, wird die Wahl der Herkunft der Pflanzen häufig nicht berücksichtigt, was teilweise darauf zurückzuführen ist, dass die Wirkung der Herkunft auf Bestäuber unbekannt ist. In dieser bahnbrechenden Fallstudie testeten wir, ob Bestäuber zwischen experimentellen Pflanzengemeinschaften verschiedener Provenienzen unterscheiden.

Wir etablierten experimentelle Pflanzengemeinschaften mit der gleichen Artenzusammensetzung, aber mit Pflanzen aus drei verschiedenen Provenienzen. Anschließend erfassten wir die Phänologie der Pflanzen und beobachteten Bestäuber und Blütenbesucher, die mit diesen experimentellen Gemeinschaften interagierten, und setzten den Besuch der Bestäuber mit der Herkunft der Pflanzen in Beziehung.

Die Provenienzen der experimentellen Pflanzengemeinschaften hatten einen starken und signifikanten Einfluss auf die Vielfalt und Häufigkeit von Blüten-Bestäuber-Interaktionen, wobei eine Provenienz doppelt so häufig interagierte wie die anderen beiden Provenienzen. Dieser Effekt wurde durch die Unterschiede in der Blühphänologie zwischen den Provenienzen hervorgerufen.

**Synthese und Anwendungen:** Pflanzenprovenienzen unterscheiden sich erheblich in ihren Interaktionen mit lokalen Bestäubern. Daher sollte die Auswahl der Pflanzenprovenienz bei der Planung von Renaturierungsprojekten zur Unterstützung von Bestäubern berücksichtigt werden.

## Introduction

As UN declared the decade 2021-2030 The Decade on Ecosystem Restoration, ecological restoration is now a political priority (www.decadeonrestoration.org). In terrestrial ecosystems, successful restoration often requires re-establishment of plant communities. These communities may establish via natural succession, but in modern, fragmented landscapes, natural succession often leads to incomplete recovery of vegetation, partly because of missing seeds in the soil or absence of natural habitats that could serve as diaspore source to recolonize the new habitat (Isbell et al. 2019). Plant introduction thus became a major tool in ecological restoration (Hölzel et al. 2012). When selecting species for introduction, there is a general agreement that restoration should prioritize native species (McDonald et al. 2016). However, there is a substantial debate on which provenance will ensure the best restoration outcome (Breed et al. 2013, 2018; Bucharova 2017; Kettenring et al. 2014; Broadhurst et al. 2008; Prober et al. 2015).

Traditionally, the local provenance was considered to be the best choice (Hamilton 2001). This is based on the observation that populations of the majority of plant species are genetically differentiated and this differentiation partly reflects adaptation to the local environment. As a result, plants of local provenance perform on average better than plants from foreign provenances, a pattern termed local adaptation (Oduor et al. 2016; Leimu & Fischer 2008; Bucharova, Durka, et al. 2017). However, with ongoing climate change, local provenancing has been increasingly criticized, because the plants are adapted to the past and current climate at the locality. As the climate changes, local adaptation is expected to lag behind (Anderson & Wadgymar 2020). Some experts thus suggest to replace or supplement the local provenance by plants from populations that are adapted to predicted climate and thus, improve performance of the restored population under climate change (Prober et al. 2015).

Throughout this provenancing debate, arguments mostly focused on plant performance. However, plants are the basis of most terrestrial ecosystems and there are myriads of other organisms that depend on them. Especially herbivores and pathogens differentiate among plant genotypes and some provenances support more herbivores than others (Macel et al. 2017; Crémieux et al. 2008; Sinclair et al. 2015; Field et al. 2019; Pearse et al. 2015). Moreover, there can be complex interaction patterns across trophic levels, so that herbivores are adapted to their local plant provenances (Runquist et al. 2019; Garrido et al. 2012; Kalske et al. 2012). And the effect of plant provenance on herbivores can even cascade up to higher trophic levels (Bucharova et al. 2016). Therefore, the choice of plant provenance for restoration is likely to affect the structure of the food webs in the restored ecosystems.

In contrast to herbivores, the effect of plant provenance on pollinators is less known. Pollination is an important ecosystem service and many restoration efforts aim to promote pollinators in the landscape (Nicholson et al. 2020; Krimmer et al. 2019; M’Gonigle et al. 2015). In this context, attention is paid to the selection of plant species that promote pollinators, but the variability within these plant species is largely ignored (M’Gonigle et al. 2017; Williams & Lonsdorf 2018; Nichols et al. 2019). The effect of provenance on pollinators was mostly demonstrated in highly specialized plant-pollinator systems, for example in orchids with one or few pollinators (Sun et al. 2014; Newman et al. 2012; Boberg et al. 2014). Such specialized plants are rarely the target of large-scale restoration, because restoration mostly relies on common foundation species that tend to be generalists (Zelený & Chytrý 2019). Nevertheless, even provenances of these common species often vary in traits that may influence their interaction with pollinators, for example in flowering phenology (Bucharova, Michalski, et al. 2017; Vander Mijnsbrugge et al. 2015; Lyngdoh et al. 2012; Díaz & Merlo 2008).

Phenology differences between provenances are genetically determined, common and documented in many species, though the exact differences between provenances is species-dependent (Vander Mijnsbrugge et al. 2015; Robson et al. 2013; Wilkinson et al. 2017; Jones et al. 2001; Bucharova, Michalski, et al. 2017). In the restoration context, selecting earlier flowering provenances would imply earlier availability of flowers and higher flower abundance early in the season. When using multiple species for restoring plant communities, the provenance of the seed mixture will affect how the flower abundance and diversity of the restored community will vary through the season, because individual plant species will start and finish their flowering depending on the species-specific provenance effects. Such differences in flower abundance and diversity are likely to modulate how the plant community interacts with pollinators, and affect pollinator identity, abundance and diversity (Fründ et al. 2010; Ebeling et al. 2008; Fornoff et al. 2017). However, the effect of plant provenance on pollinators has been only rarely studied.

In this study, we focused on the effect of plant provenance on pollinators and flower visitors. We selected six insect pollinated plant species that are an important component of species-rich seed mixtures for ecosystem restoration in Germany (Bucharova et al. 2019). For all six species, we obtained seeds from three provenances (Figure 1). We assembled small experimental plant communities that consisted of the same number of plants and the same species, but differed in provenance. We recorded plant phenology and observed pollinators and flower visitors. We hypothesized that (1) plant provenances differ in flowering phenology, which (2) results in differences in the diversity and total abundance of flowers in the experimental plant communities, and subsequently (3) affects the interacting pollinators and flower visitors, ultimately leading to differences in plant-pollinator interaction networks between provenances.

**Figure 1:**
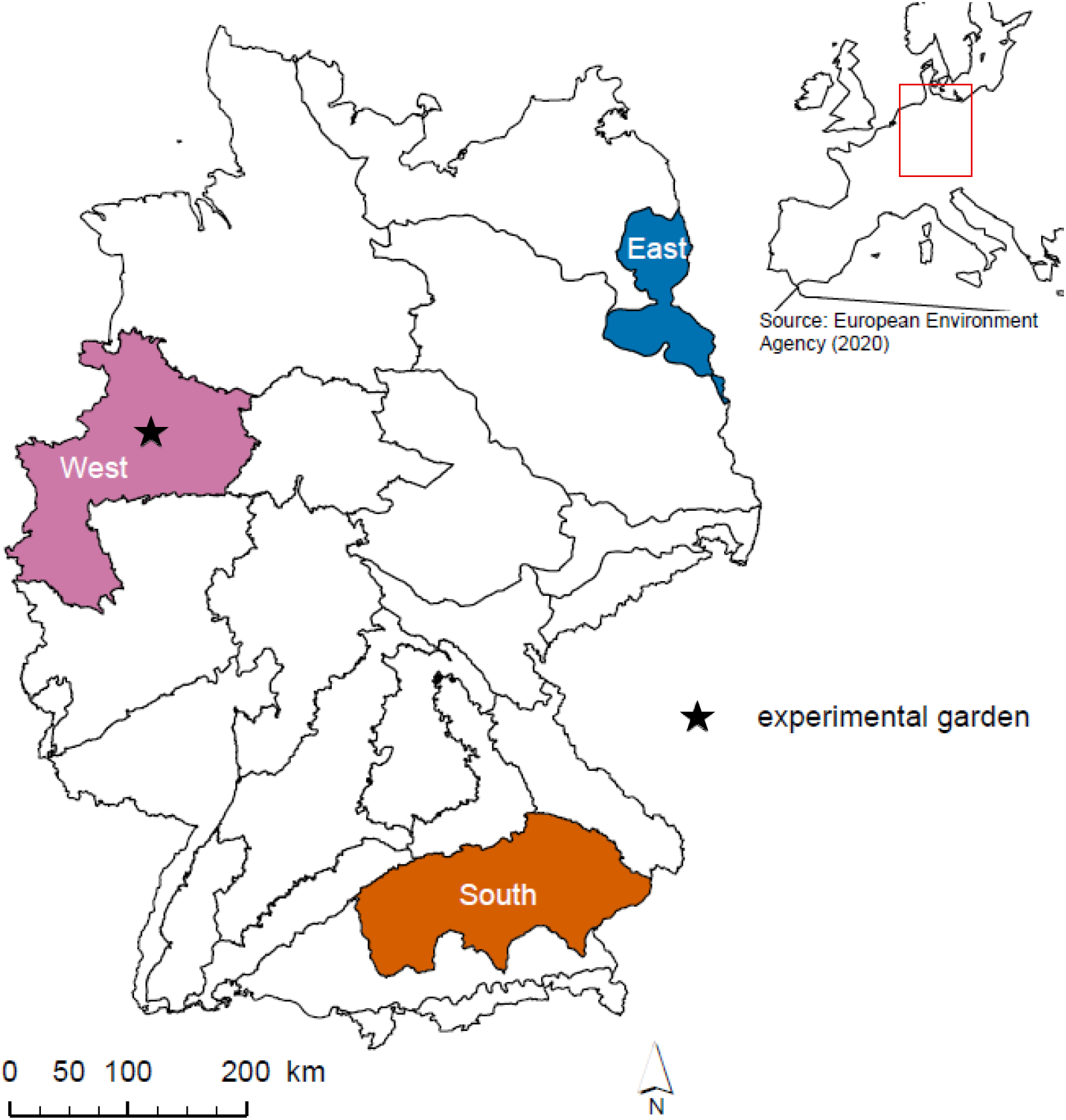
Map of seed transfer zones in Germany with highlighted provenances used in the experiment. Star indicates the location of the common garden. After Bucharova et al. (2019).

## Materials and Methods

### Selection of plants and experimental garden

We chose six insect pollinated plants species: *Centaurea jacea, Galium album, Hypochaeris radicata, Knautia arvensis, Lotus corniculatus* and *Silene vulgaris*; abbreviated by the genus name further on. All species were insect pollinated. We obtained the seeds from specialized seed producers that produce seeds for restoration according to the regional admixture provenancing (Bucharova et al. 2019; Nagel et al. 2019), Rieger-Hofmann GmbH and Saaten Zeller GmbH & Co KG. In these systems, seeds are collected from multiple populations within a given region, so called seed transfer zone, mixed and propagated on farms to produce regionally adapted, yet genetically diverse seed lots.

For all six species, we selected three provenances that span the geographic and climatic variability in Germany: a western, an eastern and a southern provenance (Figure 1). The western provenance, representing the regional provenance of our experiment, has an oceanic climate with relatively low seasonal differences, a mean annual temperature of 8-10°C and an annual precipitation of 700-800 mm. The southern provenance is slightly colder and wetter (6-8°C mean annual temperature, 700-1000 mm annual precipitation), and has a continental climate with larger differences between seasons. The eastern provenance also has a continental climate, the same mean annual temperature as the western provenance (8-10°C), but receives considerably less precipitation of 450-550 mm per year (Deutscher Wetterdienst, 2020).

We carried out this experiment in 2019 in a common garden in Münster, Germany (Figure 1). The summer of 2019 was 2.4°C warmer with only 63% of the precipitation compared with the long-term mean (Deutscher Wetterdienst, 2020). The experimental garden was located in an abandoned botanical garden, a neglected area covered by a variety of habitats ranging from open grasslands to dense shrubs. It provides abundant and diverse flower resources and also nesting opportunities for pollinators.

### Establishing experimental plant communities

We sowed seeds of all six plant species and three provenances to seeding trays in March 2019 and kept the trays in a cold greenhouse. When the seedlings developed the first true leaves, we transplanted them to planting trays. After 4-6 weeks, depending on the species, we transplanted all small plants to 1.5 l pots filled with standard potting substrate and transferred them to the common garden. Using potted plants allowed each plant to grow in standardized conditions, and avoid confounding effects of mortality due to direct competition between the plants. We arranged sets of individual potted plants (one specimen of each species) all originating from the same provenance in small patches side-by-side. In the case that any of the experimental plants died, we replaced it with a back-up plant from the same species and provenance. In this way, we established small experimental plant communities that had the very same species composition and the same plan abundances, but differed in provenance. These experimental communities were replicated nine times for each of the three provenances, resulting in 27 experimental communities and 162 plants in total (Figure S1). The position of each species in the experimental community was selected at random, as well as the arrangement of experimental communities within the experiment. The diameter of each experimental community was approximately 70 cm, and the communities were 150 cm apart. We watered the plants as needed.

Shortly after transfer to the common garden, the experiment experienced a set-back due to massive herbivory by wild rabbits. After we fenced the whole experiment, we cut all plant individuals to a height of 2cm to standardize the aboveground defoliation. All plants regenerated within a few weeks, yet, this event delayed the flowering of all plants.

### Data collection

We recorded phenology during the whole experiment. We visited the plants every week and recorded whether individual plants did flower or not.

We observed pollinators at three dates: 31 July, 14 August, and 28 August, henceforth called early, mid, and late observation. The early observation took place one week after all experimental communities produced at least one flower. At each observation date, we recorded the number of flowering units per plant in all the experimental communities. The flowering units were defined either as individual flowers (in *Silene*) or whole inflorescences (all other species), depending on the visually distinct entity in a given species. We call all flowering units “flowers” in the further text.

Subsequently, we observed flower visitors and pollinators. Not all insects that visit flowers transfer pollen, but we did not distinguish between legitimate pollinators and other visitors. However, we call all observed insects “pollinators” in the further text for the sake of simplicity. To determine pollinators, we screened pollinators present in the experimental garden in the week before the first pollinator observations. We developed a classification key that allowed us to obtain as detailed information on the taxonomic status as possible when observing the insects sitting on the flower without catchment. We divided the pollinators into six main groups: hoverflies, flies, honey bees, other wild bees, bumblebees and wasps. Some of these groups were further divided into easily recognizable morphospecies. We did not observe any butterflies or beetles. In total, we recognized 11 pollinator morphospecies (Figure 2, Figure S2).

**Figure 2:**
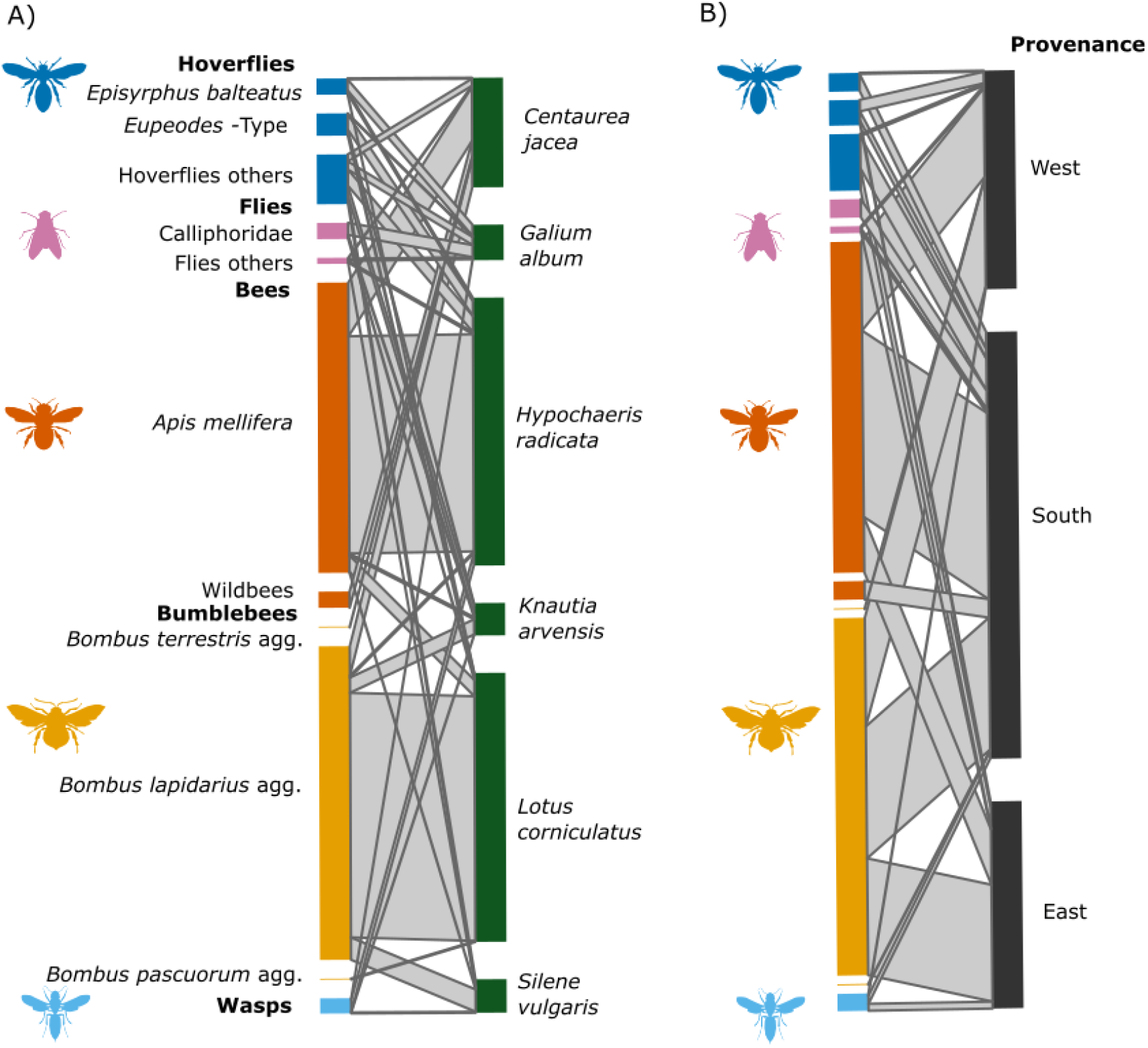
(A) All realized plant-pollinator interactions. (B) Pollinator interactions with provenance as the lower trophic level. Southern provenance attracted in total twice more pollinators than the other two provenances. The size of the boxes represents the relative abundance with which a particular pollinator or plant participated in plant-pollinator interactions.

To record pollinators, we observed each experimental community replicate for 15 minutes in two distinct 7.5 min sections, an early and a late one, to avoid the confounding effect of daytime and experimental community identity. The two sections were randomly assigned to two independent observers. All observations were conducted in stable conditions during dry weather, between 10 a.m. and 2.p.m. Within the 15 minutes, we recorded all observed flower-pollinator interactions, noting the plant species and the pollinator morphospecies. We counted individual flower-pollinator interactions, that means, when a pollinator visited several flowers at the same plant individually, we counted this as independent interactions, because the pollinator had to either fly or crawl over vegetative parts to get from one flower to another.

### Data analysis

We conducted five steps in our analyses. In the first two steps we focused on plants. We analyzed whether plants of the three provenances differed in phenology, and consequently, whether the plant provenance did affect the flower diversity of the experimental communities. In the third and fourth step we focused on the pollinators. We analyzed the plant-pollinator network and tested whether the provenance identity of the experimental plant communities effected the number and diversity of flower-pollinator interactions. In the fifth step, we connected the data on flower resources and pollinators to test whether the effect of plant provenance on pollinators was mediated by the diversity of flower resources. All these steps are detailed below. The analyses were made in R (R Development Core Team 2020).

First, we focused on plant phenology on the level of individual plants (each plant used as a replicate). As many plants did not start to flower over the course of the whole experiment, we analyzed the data on phenology in two steps, first using flowering probability with all plants, and second flowering time only with plants that did flower. For *flowering probability* during the experiment, we related flowering (binary response yes/no) to plant species, provenance and the interaction term of these two explanatory variables in a generalized linear model with a binomial error distribution. Among provenances of each individual species, we compared the probability of flowering using post-hoc pairwise chi-square comparisons. For *flowering time*, we related the onset of flowering of each individual plant to species, provenance and the interaction term of these two variables in a linear model with a gaussian error distribution. We compared provenances using pairwise comparisons as provided in the *multcomp* package (Hothorn et al. 2008). We further used the *effects* package to obtain fitted values and confidence intervals for visualization (Fox & Weisberg 2014).

Second, we focused on the provenance effect on flower abundance and flower diversity of the experimental communities, expressed by the Shannon index. As flower abundance and diversity were correlated (r=0.56), we used only flower diversity for the analyses and results presented below. The results using flower abundance largely mirrored the results using flower diversity (Table S1, Figure S3). In this and subsequent analyses, a single value per experimental community was one replicate. We used linear mixed effects models and related flower diversity per experimental community to provenance, date of observation and their interaction as fixed factors, using the identity of the experimental community as a random factor to account for repeated observations. All mixed effects models (lmm and glmm) here and later were computed with the R-package *blme* (Chung et al. 2013).

Third, we visualized all observed flower-pollinator interactions using the *bipartite* package (Dormann et al. 2008). Across all plants and species in the experiment, we used a chi-square test to test whether the number of interactions per pollinator morpho-species differed among plant provenances. Further, using a null model approach (function: *vaznull*, (Vázquez et al. 2007)), we tested if pollinators preferably visited certain provenances by comparing their observed choice of provenance to what would be expected at random. To this aim we computed the specialization index *H*_2_’ for our observed interactions and compared it with the distribution of *H*_2_’ of 1000 simulated pollinator-provenance networks (i.e. random provenance choices) with the same number of interactions. The specialization index *H*_2_’ ranges between 0 and 1 and takes on high values when a network is specialized, i.e. when the pollinator interactions were not evenly distributed among provenances.

Fourth, we analyzed the effect of provenance identity on the number of plant-pollinator interactions and the diversity of pollinators, essentially the number of pollinator taxa (a single value per experimental community was one replicate). Since the number of observations, and thus the number of plant-pollinator interactions, was often low per experimental community, we did not calculate the Shannon index but used the number of pollinator taxa (i.e., morphospecies richness) as a surrogate of insect diversity. We related this pollinator diversity and the number of interactions as response variables to provenance, observation date and their interaction term as explanatory variables in a generalized linear mixed effects model with a Poisson error distribution using experimental community as random block effect. To account for overdispersion we added an observation level random effect (i.e. with a unique level for each data point) to the model with the number of interactions as response variable (Harrison 2014).

Fifth, we connected the data on pollinators and flowers per experimental community and we tested whether flower diversity (expressed as Shannon index, see above) affects the number of plant-pollinator interactions and the pollinator diversity (i.e., the number of pollinator taxa, see above). We related the pollinator diversity and the number of plant-pollinator interactions as response variables to flower diversity, date of observation and their interaction and provenance as explanatory variables in a generalized linear mixed model with a Poisson error distribution and the experimental community as random block effect. In a second step, we added provenance as an explanatory variable to this model, to analyze if provenance had an additional effect on top of the flower diversity effect (i.e. if provenance explained any variability when fitted after flower diversity and date), or if the provenance effect was driven by the flower diversity effect. Again, an observation level effect was used to account for overdispersion (Harrison 2014).

## Results

### Plant phenology and flower diversity

Plant provenance affected both probability of flowering during the experiment and flowering time (Figure 3, Table 1A, Figure S4). This effect was especially strong in two species, *Galium* and *Centaurea*, with the southern provenance flowering more frequently than the other two provenances (Figure 3A, B). In these species, the southern provenance also flowered relatively early in comparison to the western provenance (Figure 3B, Table 1A). In *Hypochaeris*, flowering time followed a similar trend, but the effects were not significant (Figure 3A, Table 1A).

**Table 1:** Results of (A) the model testing the effect of provenance, species and their interactions on plant phenology. (B) Model testing the effect of provenance, observation date and their interaction on flower diversity, pollinator diversity and number of flower-pollinator interactions. (C) Model testing the effect of flower diversity, observation date and their interaction on pollinator diversity and number of flower-pollinator interactions. Provenance was added to the model (C) after testing for significance of the other terms. All models are with error type III; significant values are in bold.

|  | Df | Flowering probability (glm) |  | Onset of flowering (lm) |  |  |
| --- | --- | --- | --- | --- | --- | --- |
|  |  | X<sup>2</sup> | P | Resid DF | F | P |
| Species | 5 | 44.9 | **<0.001** | 94 | 19.96 | **<0.001** |
| Provenance | 2 | 19.65 | **<0.001** | 94 | 8.76 | **<0.001** |
| Species x Provenance | 10 | 13.62 | 0.191 | 94 | 2.81 | **0.004** |

B) Flower and insect diversity
|  | Df | Flowers (lmm) |  | Pollinators (glmm) |  |  |  |
| --- | --- | --- | --- | --- | --- | --- | --- |
|  |  | Diversity |  | Number of interactions |  | Pollinator diversity |  |
|  |  | X<sup>2</sup> | P | X<sup>2</sup> | P | X<sup>2</sup> | P |
| Provenance | 2 | 45.07 | **<0.001** | 12.41 | **0.001** | 12.61 | **0.002** |
| Observation date | 2 | 10.11 | **0.006** | 0.54 | 0.764 | 14.00 | **<0.001** |
| Provenance x Observation date | 4 | 7.35 | 0.118 | 6.63 | 0.156 | 12.35 | **0.015** |

C) Effect of flower diversity on pollinators
|  | Df | Pollinator diversity (glmm) |  | Number of interactions (glmm) |  |
| --- | --- | --- | --- | --- | --- |
|  |  | X<sup>2</sup> | P | X<sup>2</sup> | P |
| Flower diversity | 1 | 9.37 | **0.002** | 7.48 | **0.006** |
| Observation date | 2 | 6.43 | **0.040** | 8.16 | **0.017** |
| Flower diversity x Observation date | 2 | 7.40 | **0.025** | 6.27 | **0.043** |
| Provenance | 2 | 1.08 | 0.583 | 0.061 | 0.970 |

**Figure 3:**
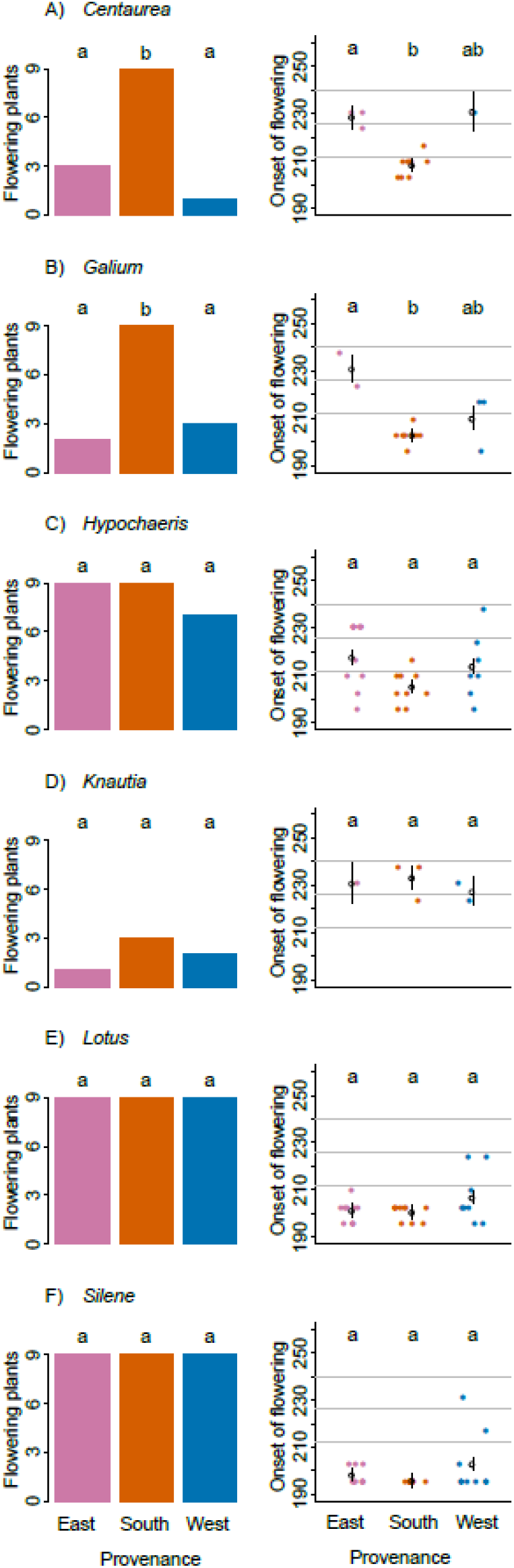
Probability of flowering during the experiment (left column) and the onset of flowering for plants that started to flower (right column), for the six species included in the experiment. Letters mark significant differences between provenances. Grey horizontal lines in the right column indicate dates of pollinator observations. For the results of overall models, see Table 1A.

The phenological differences resulted in a different flower diversity among provenances at a given date, with the experimental communities of southern provenance showing a two-fold higher flower diversity than the experimental communities of the other two provenances. This effect was consistent across observation dates (Figure 4A, B, Table 1B).

**Figure 4:**
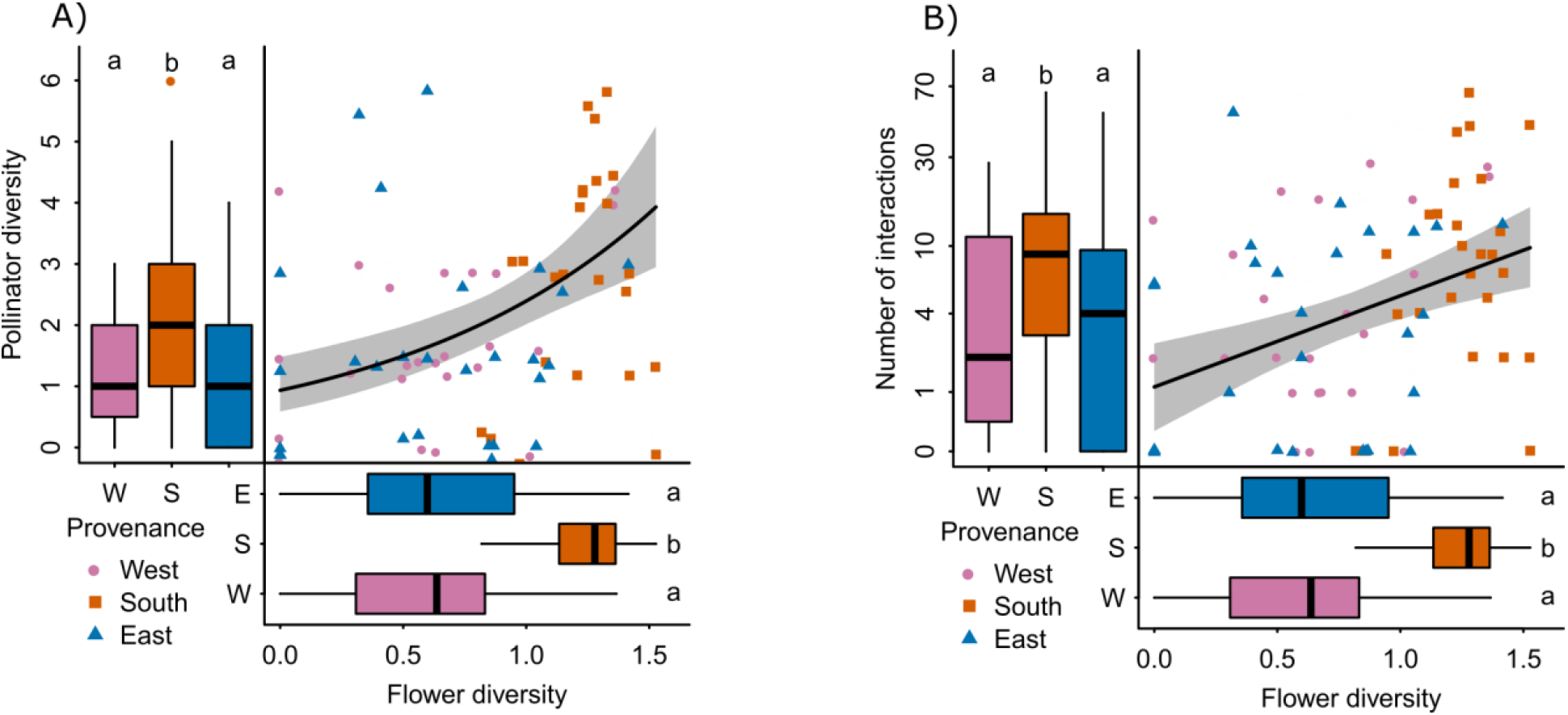
The relationship between flower diversity and (A) pollinator diversity, expressed as the number of taxa, and (B) pollinator abundance. Horizontal boxplots show the provenance effect on flower diversity, the same for A and B. Vertical boxplots show provenance effect on (A) pollinator diversity and (B) number of interactions. Note logarithmic y-axis in (B). Letters indicate significant differences. For model results, see Table 1B,C.

### Pollinators

In total, we observed 737 flower-pollinator interactions. The majority (> 85%) of all interactions took place with three plant species only: *Lotus, Hypochaeris*, and *Centaurea* (265, 264 and 108 interactions, respectively). In contrast, flowers of the other three species, namely *Galium, Silene* and *Knautia*, were visited much less frequent (35, 33 and 32 interactions, respectively). The two most common insects with the majority of interactions were bumblebees (morphospecies *Bombus lapidarius*, L 1758; 309 interactions), and honey bees (*Apis mellifera*, L 1758; 286 interactions). Bumblebees preferably interacted with *Lotus*, whereas honey bees preferentially interacted with both species from the family Asteraceae – *Centaurea* and *Hypochaeris* (Figure 2A).

Across all plants and species, plants of the southern provenance interacted with pollinators twice as often as the eastern and western provenance (yielding 369, 189 and 189, respectively; Figure 2B). Notably, while in the eastern provenance bumblebees accounted for 60% of all interactions, their share was lower in the western and southern provenance (Figure 2C). This indicates that pollinators differentiated between the provenances, which was further suported by a moderate specialization index *H*_2_’ (*H*_2_’= 0.1) for the network calculated across all monitoring dates. The index was significantly higher than zero (P < 0.001, Figure S5), which means that insects visited one of the provenances more often than would be expected if the visits were random (Figure 2B).

Accordingly, the provenance identity of the plant experimental communities affected the diversity of pollinators (expressed as the number of pollinator taxa), with the southern provenance interacting with more taxa than the western or eastern provenance (Figure 4A, Table 1B). The number of interactions was on average also the highest in the experimental communities of the southern provenance, but this effect depended on the date of observation (Figure 4B, Table 1B, Figure S6). Experimental communities with higher flower diversity interacted with more pollinator taxa and had a higher number of interactions. (Figure 4 A, B, Table 1C). The slope of these relationships differed between observation dates, but overall, the effect remained positive (Figure 4 A, B, Table 1C, Figure S4). When we added provenance to this model, it had no significant effect, indicating that the provenance effect was mediated by flower diversity.

## Discussion

We tested the effect of within-species population differentiation on the pollinator diversity and the frequency of flower visitation by pollinators and flower visitors (further pollinators for simplicity) in experimental plant communities that consisted of the very same plant species, but differed in their provenance identity. We then recorded the plant phenology, the number of flowers and observed the pollinators. We showed that plant phenology differed across plant provenance identities, a pattern mediated by differences in flower diversity among provenances. In consequence, the identity of plant provenance had a profound effect on the frequency of flower-pollinator interactions as well as on pollinator diversity. Amongst plant provenances, the southern identity interacted twice as often with pollinators and attracted twice as many species as each of the other two provenances.

### Plant phenology

Plants from the three provenances differed in phenology. Plants coming from the southern provenance (continental climate) flowered earlier and more often than plants from the other two provenances, especially, in comparison to the western provenance, which has an oceanic climate. This effect was significant in two out of six species. Such differences in phenology of provenances are common among plants. Bud burst, leaf flush, flowering phenology or senescence are important parts of plant adaptation to their environment and they are partly genetically underpinned (Vitasse et al. 2009; Vander Mijnsbrugge et al. 2015; Cooper et al. 2019; Zhang et al. 2019; Bucharova, Michalski, et al. 2017). Our results partly correspond to the fact that in temperate regions, plants from warmer and more continental origins typically show advanced phenology when compared to plants from northern origins and more oceanic climates (Vander Mijnsbrugge et al. 2015; Robson et al. 2013; Wilkinson et al. 2017; Jones et al. 2001).

However, attributing the phenology differences to climate at origin must be done with caution. We have watered the plants throughout the experiment, and it is possible that the provenance that has profited the most from the surplus of water started to flower earlier. Further, we only observed the phenology in the first year after germination in perennial plants, ignoring the possibility that the phenology in the year of establishment may differ from that in later years. Yet, phenology differences among provenances are highly genetically determined and they tend to be similar across the years (Liang 2015; Jones et al. 2001; Smith et al. 2020). Further, the experiment also suffered from massive herbivory damage by wild rabbits, which almost completely removed aboveground biomass. The plants thus had to first regrow, and the observed difference in flowering could be caused by a faster regrowth of some provenances compared to others. Yet, this is rather unlikely: First, similar phenology differences among provenances as in the current study, but without herbivory damage, were observed in a previous study with a larger set of plants from the same ecosystem (Bucharova, Michalski, et al. 2017). Secondly, all the plants are characteristic species of human-managed grasslands, they are adapted to regular mowing and rapid re-growth is part of their survival strategy (Bucharova et al. 2020).

On the community level, the differences in flower phenology among provenances caused that experimental communities of the later flowering provenances lacked flowers of the species with delayed phenology (i.e., *Centaurea* and *Gallium*), especially during the first observation dates. This necessarily resulted in differences in flower abundance and diversity among provenances, with the early flowering southern provenance having on average a more than twofold higher flower diversity.

### Pollinators and flower visitors

The differences among provenances in flower diversity resulted in differences in the number and diversity of flower-pollinator interactions. Specifically, experimental communities of plants from the earlier flowering provenance (i.e. southern) had higher flower diversity and consequently, their flowers interacted more frequently with pollinators and flower visitors, and the interacting insects were more diverse. Flower diversity is, in fact, a well-known predictor of pollinator diversity (Fründ et al. 2010; Ebeling et al. 2008; Fornoff et al. 2017). Yet, the evidence so far came from observations of communities that differed in plant species richness. Since we kept the species diversity constant by harmonizing species identities and richness across provenances, we illustrate how important is the variability within plant species for the interaction with pollinators.

The effect of plant provenance on the diversity of interacting pollinators is likely underestimated in our study. We differentiated pollinators only to 11 morphospecies that were easy to identify when observing insects sitting on the flowers. These morphospecies likely comprise several true species that differ in seasonality. It is thus possible that for example hoverflies or wild bees interacting with flowers during the first observation belonged to a different species than individuals of the same morphospecies observed at the last observation date (Ball & Morris 2015 Westrich (2018) for wild bees). Our estimate of provenance effect on pollinators is thus rather conservative.

### Implications for ecosystem restoration

We have demonstrated that the provenance identity of plant experimental communities alters the diversity of flowers of these experimental communities which subsequently affects the frequency of flower-pollinator interactions and the diversity of interacting insects. The provenance selection for ecosystem restoration is intensively debated, especially in the context of ongoing climate change (Breed et al. 2013, 2018; Prober et al. 2015), but the arguments are mostly about plant performance. As ecological restoration aims to restore all components of ecosystems, ecosystem functions and services, the effect of provenance choice on interacting biota including herbivores and pollinators should not be neglected. This is especially the case in projects that aim on supporting pollinators in agricultural landscapes (Nicholson et al. 2020; Krimmer et al. 2019; M’Gonigle et al. 2015). In this context, attention is usually paid to the selection of particular plant species that are beneficial to pollinators, but within-species variability of these plants is largely neglected (M’Gonigle et al. 2017; Williams & Lonsdorf 2018; Nichols et al. 2019). We show that within-species variability may have a substantial impact on how flower resources are available throughout the season, potentially leading to a decoupling of plant-pollinator interactions. Such decoupling may shift the abundance and diversity of pollinator interactions, as generalist pollinators that collect pollen from multiple species will likely be less affected than species specialized on one or a few plants (Visser & Gienapp 2019; Renner & Zohner 2018). In consequence, the properties of the plant-pollinator interaction network will likely depend on the phenology of the provenance that was used for a given restoration project (Bartley et al. 2019).

Caution is needed when interpreting our results on the landscape level because of methodology. We used an experimental set up in which we offered experimental communities of different provenance to pollinators within one garden and the pollinators were given the choice. In reality, restored habitats are usually seeded with one provenance and pollinators cannot choose. Yet, the selection of provenance will affect the availability of flower resources, which will subsequently influence the abundance and diversity of pollinators in the whole restored habitat (Fründ et al. 2010). Further, we used potted plants to establish the experimental communities, which reduced competition between plants. In restoration, plants are seeded or planted and they compete for resources. The competitive ability of individual plants species differs between provenances, but the provenance effect is not consistent across species (Bischoff et al. 2006). Competition at a restoration site thus could suppress or promote different species depending on the specific provenance, which would further increase differentiation between the communities (Weißhuhn et al. 2012). Consequently, in real restoration projects, the impact of the provenance selection on pollinators is potentially larger than shown in this study.

The highest pollinator diversity did not support the local provenance, but a provenance from a different region instead. Shall we plant foreign provenances to support pollinators? The pollinators observed in our study were mostly generalists and thus, they profited from the earlier start of the flowering period (Iler et al. 2013). More specialized pollinators, however, may suffer because they are more sensitive to temporal decoupling from their forage plants (Kudo & Ida 2013). Besides the consequences for pollinator interactions, plant phenology is also an important part of adaptation to local climatic conditions (Robson et al. 2013; Cooper et al. 2019), and the foreign provenances that differ in phenology may have reduced fitness because of, for example, frost damage.

In summary, our study provides proof of principle that provenance selection does affect pollinator abundance and diversity in the restored habitat. Yet, more research is needed to formulate conclusive recommendations on the choice of the optimal provenance. Specifically, we need more data on the adaptation of plants to their environment, on co-adaptation between the plants and local pollinators, optimally obtained from reciprocal transplant experiments (Bucharova, Durka, et al. 2017).

## Acknowledgements

This work was supported by LA4038/2-1 to AB. We are grateful to Norbert Hölzel, Christoph Scherber and Theresa Klein-Raufhake for support.

## Authors’ contribution

AB conceived the idea; AB, DO, EM, MM and JM designed the study; EM, MC and JM conducted the experiment; AB and CL analyzed the data; AB led the writing of the manuscript. All authors contributed critically to the drafts and gave final approval for publication.

## Data availability statement

Data available via the Dryad Digital Repository https://doi.org/10.5061/dryad.m905qfv0x(Bucharova et al, 2021).

## Supplementary material

**Table S1:**
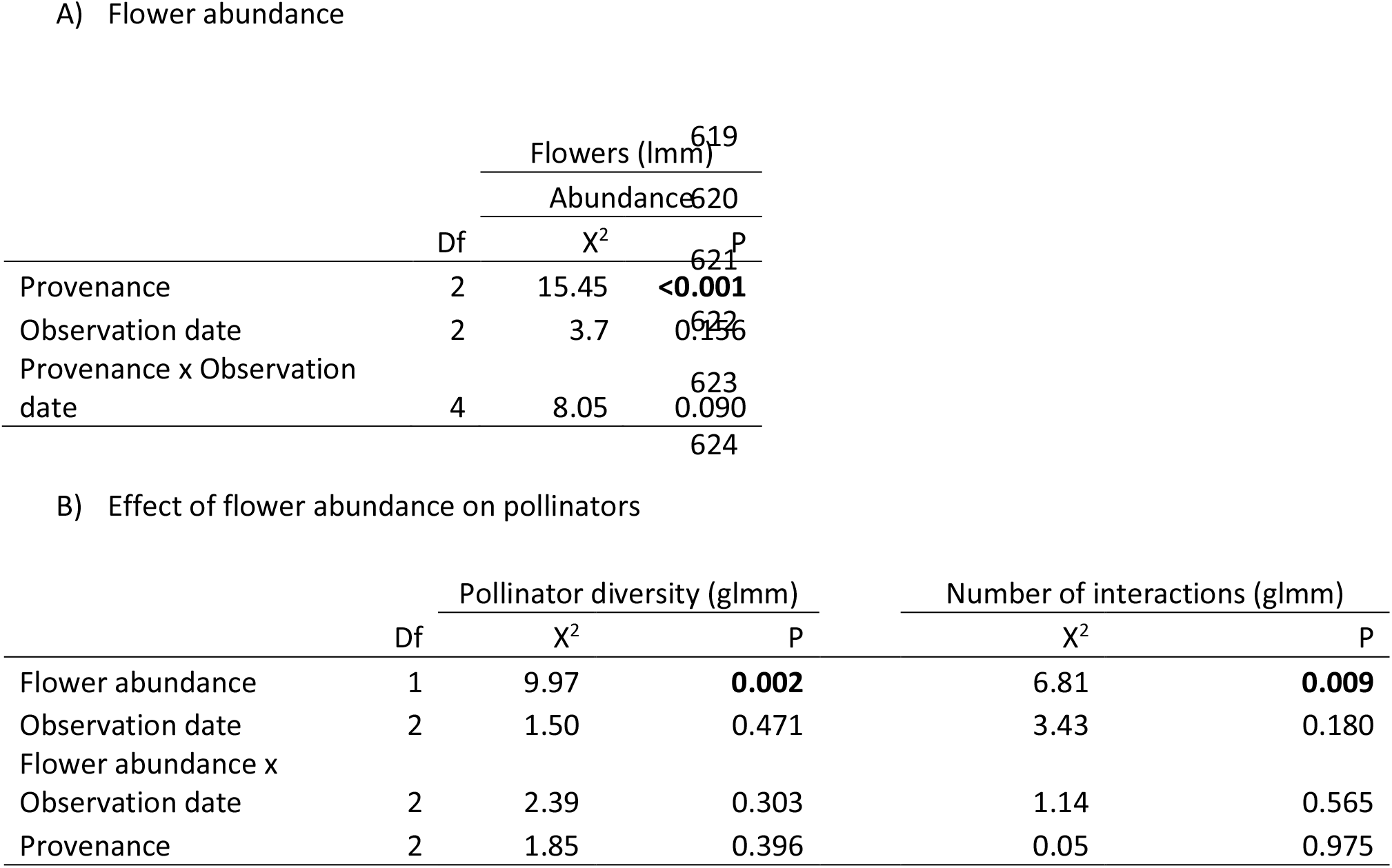
Results of the models testing (A) the effect of provenance and date of observation on flower abundance and (B) the effect of flower abundance, observation date and their interaction on pollinator diversity and number of flower-pollinator interactions. Provenance was added to the model (B) after testing for significance of the other terms. All models are with error type III; significant values are in bold.

**Figure S1:**
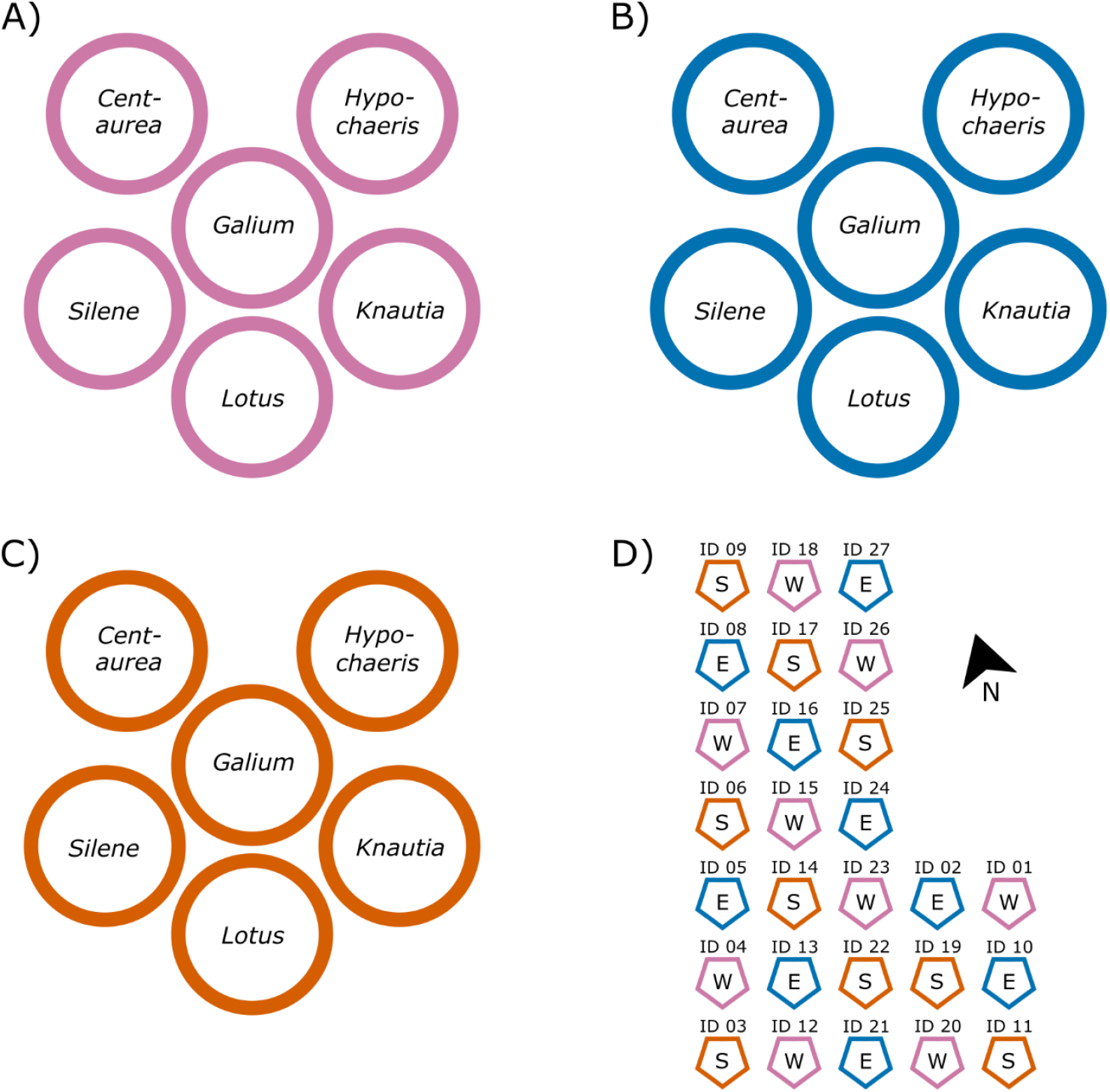
Experimental design. We selected six insect pollinated species (i.e. *Centaurea jacea, Galium album, Hypochaeris radicata, Knautia arvensis, Lotus corniculatus* and *Silene vulgaris*), and used seeds for each plant species from three different regional pools (i.e. provenances). These species were arranged in experimental communities according to their provenance, with individual potted plants, all consisting of the same species mix. Thus, we established (A) a western (origin of study side) provenance (B) an eastern provenance and (C) a southern provenance. Each provenance or experimental community was replicated 9 times. These communities were arranged in the botanical garden in a set of modified Latin squares (D), with randomized positioning of the position of the potted species in the communities. The different provenances are indicated by corresponding letters (W= western, E=eastern and S=southern provenance). In total, we established 162 potted plant individuals in 27 communities). The diameter of one experimental community was 70 cm, the distance between communities 1.5 m.

**Figure S2:**
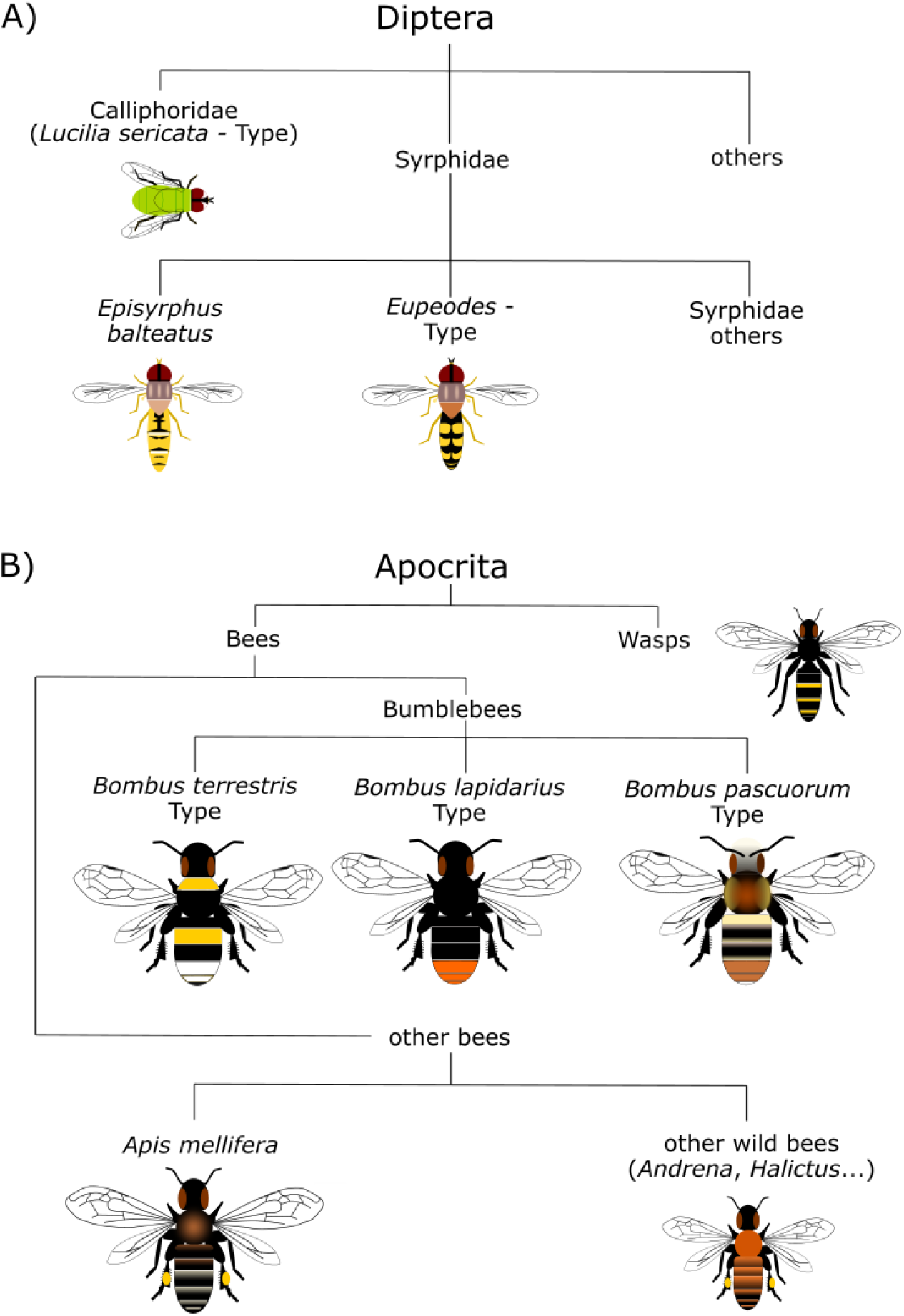
Pollinator identification. A) Amongst hoverflies (Syrphidae), that are easily to identify according to their typical flight mode, we distinguished between Episyrphus balteatus (typical moustache-pattern on the back) and Eupeodes-Type, which have a color-pattern typical for many species of the Syrphini, and other Syrphidae (not further identified). From other Dipterans we only put Calliphoridae of *Lucilia sericata*-Type as separate group. All other flies were not distinguished due to low individual counts per morphospecies. B) Bumblebees were identified according to their shape and body size. We classified them into three groups according to common and widespread species in central Europe (Esser et al., 2009; Westrich, 2018): *Bombus lapidarius-*Type (black with orange tip of abdomen), *Bombus pascuorum*-Type (blackish but with orange tip of abdomen and thorax) and *Bombus terrestris*-Type (blackish with white tip of abdomen and a yellow segment at that proximate end of abdomen and thorax). Honeybees (*Apis mellifera*) were identified to species level according to size, color and the elongated “Radial-cell” of the first wing. Other bees (usually smaller but with different wing morphology) were grouped as Wild bees and not further distinguished. We treated wasps as one group because identification to species level was not possible in the field and the individual richness of this group was in general very low.

---

**Figure S3:**
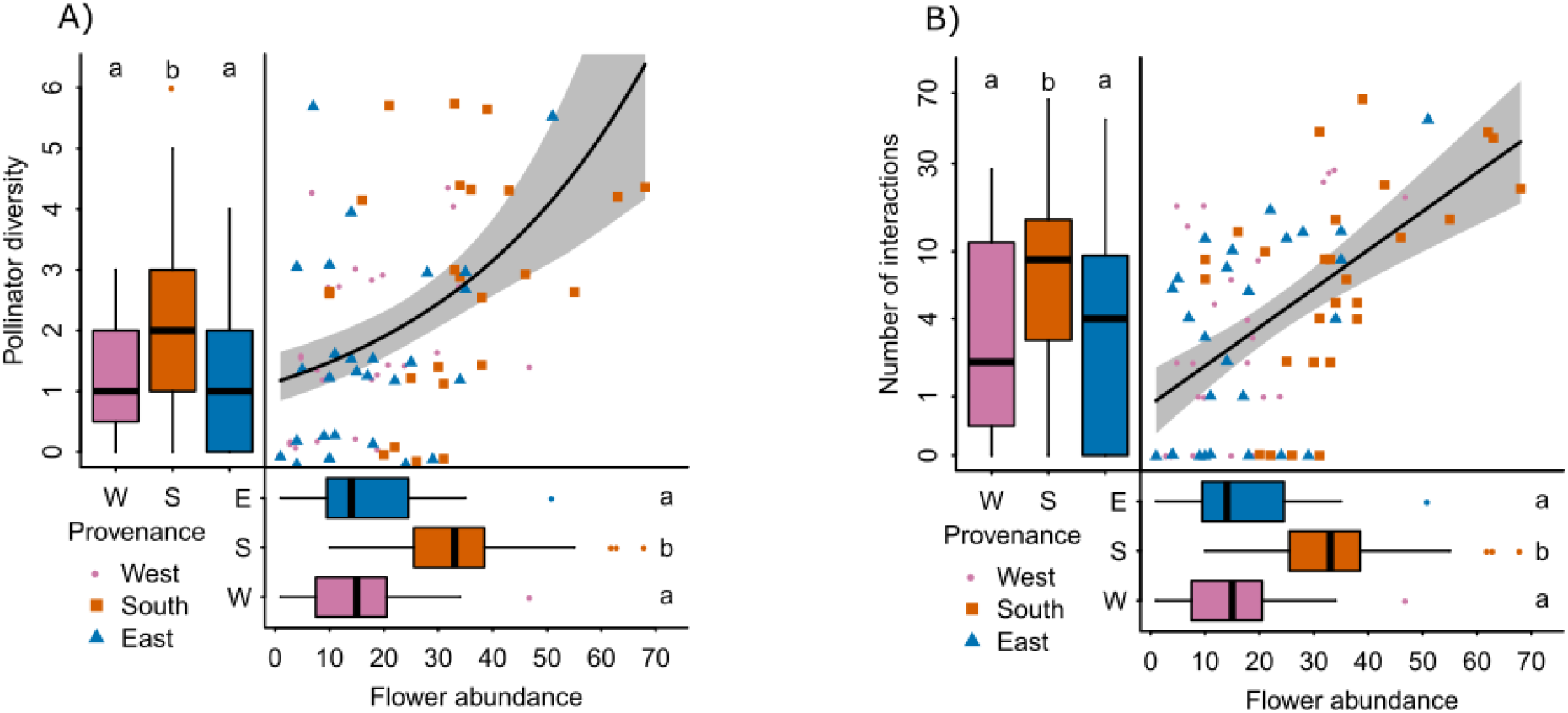
The relationship between flower abundance and (A) pollinator diversity, expressed as the number of taxa, and (B) pollinator abundance. Horizontal boxplots show the provenance effect on flower abundance, the same for A and B. Vertical boxplots show provenance effect on (A) pollinator diversity and (B) number of interaction. Note logarithmic y-axis in (B). Letters indicate significant differences. For model results, see Table 1B and Table S1A, B.

**Figure S4:**
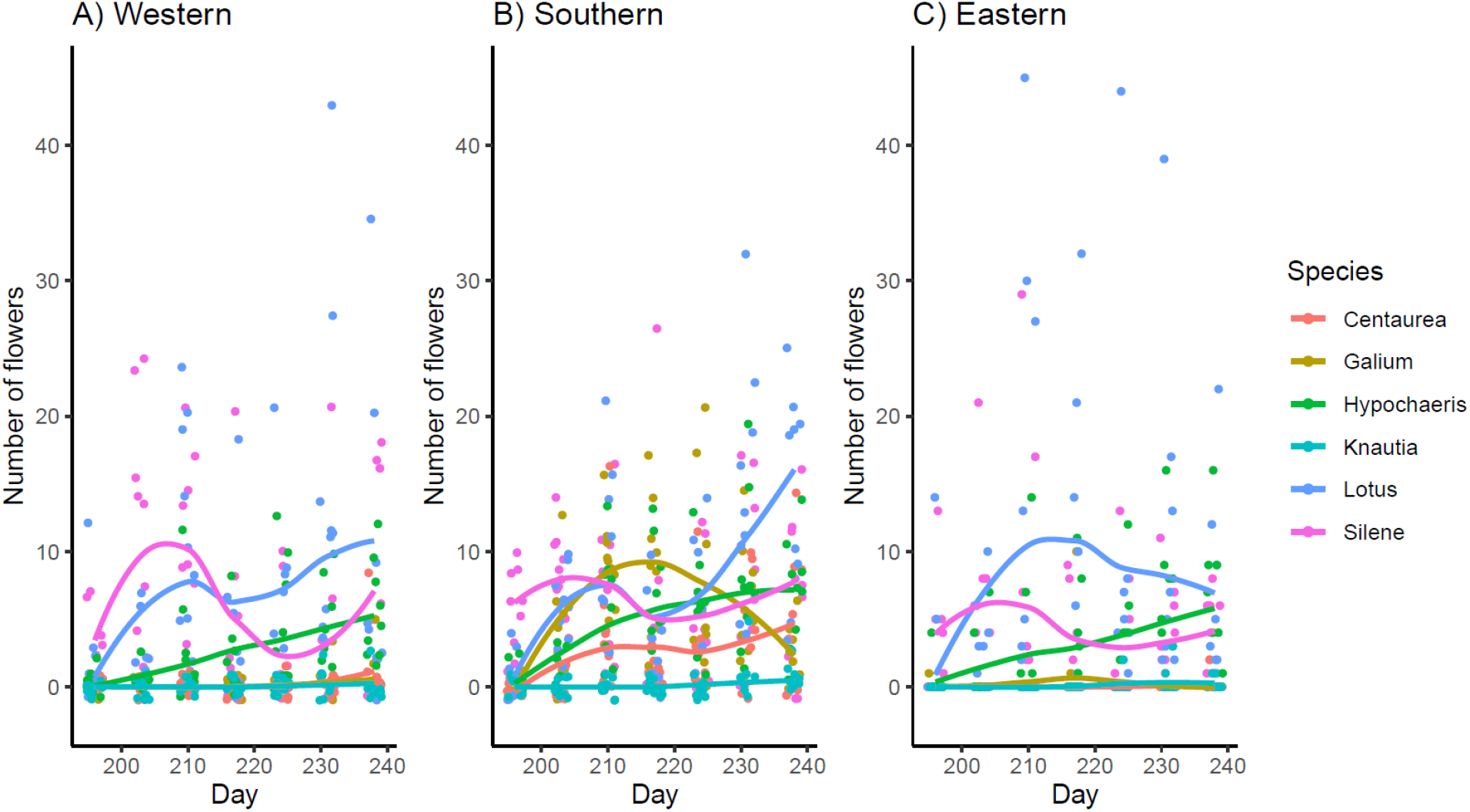
Number of flowers per each species and experimental community during the time for experimental communities of (A) western, (B)southern and (C) eastern provenance. Each point the number of flowers of given species in one experimental community in a given day. The line is smoothed line using function *loess* as implemented in *ggplot2* package.

**Figure S5.**
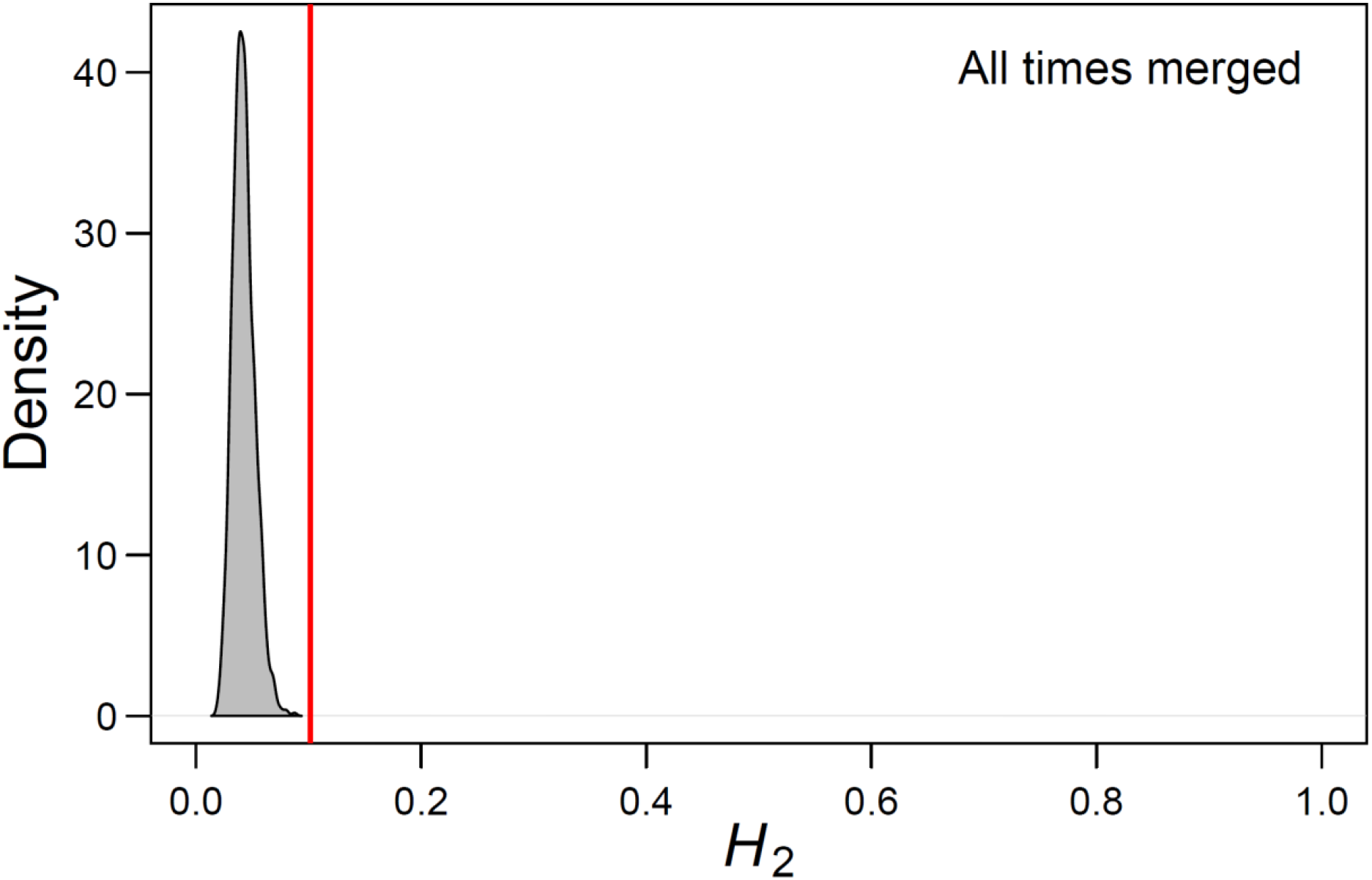
Specialization displayed as index *H*_2_’. The red vertical line indicates the observed *H*_2_’ for the network displayed in Fig. 3B. The distribution of *H*_2_’ obtained from 1000 null models with the same number of interactions and species/provenances (*bipartite* package in R, function: *vaznull*, Vázques et al. 2007). This indicates that the insect species were more specific in their interactions than would be expected at random

**Figure S6:**
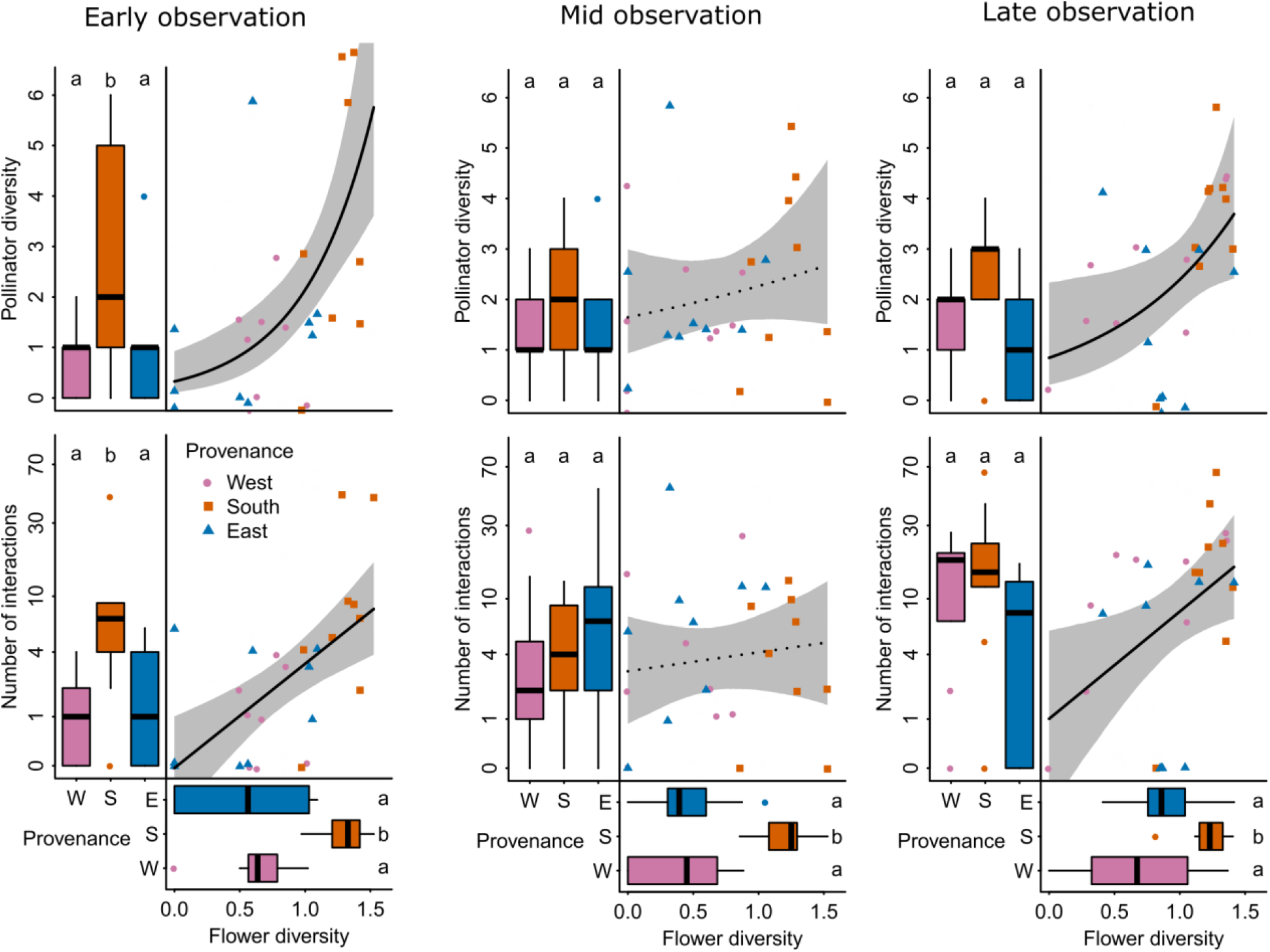
The relationships between flower diversity, pollinator diversity and number of interactions, broken into three observation dates. Upper graphs represent pollinator diversity, expressed and the number of taxa, middle graphs number of interactions, and how these variables are affected by provenance (boxplots) and flower diversity (scatterplot). Horizontal boxplots show the provenance effect on flower diversity. Note logarithmic y-axis in pollinator abundance. Letters indicate significant differences.

## References

Anderson, J. T., & Wadgymar, S. M. (2020). Climate change disrupts local adaptation and favours upslope migration. Ecology Letters, 23(1), 181–192.

Bischoff, A., Crémieux, L., Smilauerova, M., Lawson, C. S., Mortimer, S. R., Dolezal, J., … Müller-Schärer, H. (2006). Detecting local adaptation in widespread grassland species - the importance of scale and local plant community. Journal of Ecology, 94(6), 1130–1142.

Boberg, E., Alexandersson, R., Jonsson, M., Maad, J., Ågren, J., & Nilsson, L. A. (2014). Pollinator shifts and the evolution of spur length in the moth-pollinated orchid Platanthera bifolia. Annals of Botany, 113(2), 267–275.

Breed, M. F., Harrison, P. A., Bischoff, A., Durruty, P., Gellie, N. J. C., Gonzales, E. K., … Bucharova, A. (2018). Priority actions to improve provenance decision making. BioScience, 68(7), 510–516.

Breed, M. F., Stead, M. G., Ottewell, K. M., Gardner, M. G., & Lowe, A. J. (2013). Which provenance and where? Seed sourcing strategies for revegetation in a changing environment. Conservation Genetics, 14(1), 1–10.

Broadhurst, L. M., Lowe, A., Coates, D. J., Cunningham, S. A., McDonald, M., Vesk, P. A., & Yates, C. (2008). Seed supply for broadscale restoration: maximizing evolutionary potential. Evolutionary Applications, 1(4), 587–597.

Bucharova, A. (2017). Assisted migration within species range ignores biotic interactions and lacks evidence. Restoration Ecology, 25(1), 14–18.

Bucharova, A., Bossdorf, O., Hölzel, N., Kollmann, J., Prasse, R., & Durka, W. (2019). Mix and match: regional admixture provenancing strikes a balance among different seed-sourcing strategies for ecological restoration. Conservation Genetics, 20(1), 7–17.

Bucharova, A., Durka, W., Hölzel, N., Kollmann, J., Michalski, S., & Bossdorf, O. (2017). Are local plants the best for ecosystem restoration? It depends on how you analyze the data. Ecology and Evolution, 7(24), 10683–10689.

Bucharova, A., Frenzel, M., Mody, K., Parepa, M., Durka, W., & Bossdorf, O. (2016). Plant ecotype affects interacting organisms across multiple trophic levels. Basic and Applied Ecology, 17(8), 688– 695.

Bucharova, A., Lampei, C., Conrady, M., Matheja, J., Meyer M., Ott, D. (2021): Data from: Plant provenance affects pollinator network: implications for ecological restoration. ?Dryad Digital Repository, https://doi.org/10.5061/dryad.m905qfv0x

Bucharova, A., Michalski, S., Hermann, J.-M., Heveling, K., Durka, W., Hölzel, N., … Bossdorf, O. (2017). Genetic differentiation and regional adaptation among seed origins used for grassland restoration: lessons from a multi-species transplant experiment. Journal of Applied Ecology, 54(1), 127–136.

Chung, Y., Rabe-Hesketh, S., Dorie, V., Gelman, A., & Liu, J. (2013). A nondegenerate penalized likelihood estimator for variance parameters in multilevel models. Psychometrika, 78(4), 685–709.

Crémieux, L., Bischoff, A., Smilauerová, M., Lawson, C. S., Mortimer, S. R., Dolezal, J., … Steinger, T. (2008). Potential contribution of natural enemies to patterns of local adaptation in plants. The New Phytologist, 180(2), 524–533.

Díaz, R., & Merlo, E. (2008). Genetic variation in reproductive traits in a clonal seed orchard of Prunus avium in northern Spain. Silvae Genetica, 57(3), 110–118.

Dormann, C. F., Gruber, B., & Fründ, J. (2008). Introducing the bipartite package: Analysing ecological networks. R News, 8(2), 8–11.

Field, E., Schönrogge, K., Barsoum, N., Hector, A., & Gibbs, M. (2019). Individual tree traits shape insect and disease damage on oak in a climate-matching tree diversity experiment. Ecology and Evolution, 9(15), 8524–8540.

Fox, J., & Weisberg, S. (2014). An R Companion to Applied Regression: Appendices. Robust Regression in R, (December), 1–17.

Garrido, E., Andraca-Gómez, G., & Fornoni, J. (2012). Local adaptation: Simultaneously considering herbivores and their host plants. New Phytologist, 193, 445–453.

Hamilton, N. R. S. (2001). Is local provenance important in habitat creation? A reply. Journal of Applied Ecology, 38(6), 1374–1376.

Harrison, X. A. (2014). Using observation-level randomeffects to model overdispersion in count data in ecology and evolution. PeerJ, 2014(1), e616.

Hölzel, N., Buisson, E., & Dutoit, T. (2012). Species introduction - a major topic in vegetation restoration. Applied Vegetation Science, 15(2), 161–165.

Hothorn, T., Bretz, F., & Westfall, P. (2008). Simultaneous inference in general parametric models. Biometrical Journal, 50(3), 346–363.

Isbell, F., Tilman, D., Reich, P. B., & Clark, A. T. (2019). Deficits of biodiversity and productivity linger a century after agricultural abandonment. Nature Ecology and Evolution, 3(11), 1533–1538.

Jones, A. T., Hayes, M. J., & Sackville Hamilton, N. R. (2001). The effect of provenance on the performance of Crataegus monogyna in hedges. Journal of Applied Ecology, 38(5), 952–962.

Kalske, A., Muola, A., Laukkanen, L., Mutikainen, P., & Leimu, R. (2012). Variation and constraints of local adaptation of a long-lived plant, its pollinators and specialist herbivores. Journal of Ecology, 100(6), 1359–1372.

Kettenring, K. M., Mercer, K. L., Reinhardt Adams, C., & Hines, J. (2014). Application of genetic diversity-ecosystem function research to ecological restoration. Journal of Applied Ecology, 51(2), 339–348.

Krimmer, E., Martin, E. A., Krauss, J., Holzschuh, A., & Steffan-Dewenter, I. (2019). Size, age and surrounding semi-natural habitats modulate the effectiveness of flower-rich agri-environment schemes to promote pollinator visitation in crop fields. Agriculture, Ecosystems and Environment, 284.

Leimu, R., & Fischer, M. (2008). A meta-analysis of local adaptation in plants. PloS One, 3(12), e4010.

Lyngdoh, N., Gunaga, R. P., Joshi, G., Vasudeva, R., Ravikanth, G., & Shaanker, R. U. (2012). Influence of geographic distance and genetic dissimilarity among clones on flowering synchrony in a Teak (Tectona grandis Linn. f) clonal seed orchard. Silvae Genetica, 61(1–2), 10–18.

M’Gonigle, L. K., Ponisio, L. C., Cutler, K., & Kremen, C. (2015). Habitat restoration promotes pollinator persistence and colonization in intensively managed agriculture. Ecological Applications, 25(6), 1557–1565.

M’Gonigle, L. K., Williams, N. M., Lonsdorf, E., & Kremen, C. (2017). A tool for selecting plants when restoring habitat for pollinators. Conservation Letters, Vol. 10, pp. 105–111.

Macel, M., Dostálek, T., Esch, S., Bucharová, A., van Dam, N. M., Tielbörger, K., … Münzbergová, Z. (2017). Evolutionary responses to climate change in a range expanding plant. Oecologia, 184(2), 543– 554.

McDonald, T., Gann, G. D., Jonson, J., & Dixon, K. W. (2016). International standards for the practice of ecological restoration - including principles and key concepts. Washington, D.C.: Society for Ecological Restoration.

Nagel, R., Durka, W., Bossdorf, O., & Bucharova, A. (2019). Rapid evolution in native plants cultivated for ecological restoration: not a general pattern. Plant Biology, 21(3), 551–558.

Newman, E., Anderson, B., & Johnson, S. D. (2012). Flower colour adaptation in a mimetic orchid. Proceedings of the Royal Society B: Biological Sciences, 279(1737), 2309–2313.

Nichols, R. N., Goulson, D., & Holland, J. M. (2019). The best wildflowers for wild bees. Journal of Insect Conservation, 23(5–6), 819–830.

Nicholson, C. C., Ward, K. L., Williams, N. M., Isaacs, R., Mason, K. S., Wilson, J. K., … Ricketts, T. H. (2020). Mismatched outcomes for biodiversity and ecosystem services: testing the responses of crop pollinators and wild bee biodiversity to habitat enhancement. Ecology Letters, 23(2), 326–335.

Oduor, A. M. O., Leimu, R., & van Kleunen, M. (2016). Invasive plant species are locally adapted just as frequently and at least as strongly as native plant species. Journal of Ecology, 104(4), 957–968.

Pearse, I. S., Baty, J. H., Herrmann, D., Sage, R., & Koenig, W. D. (2015). Leaf phenology mediates provenance differences in herbivore populations on valley oaks in a common garden. Ecological Entomology, 40(5), 525–531.

Prober, S. M., Byrne, M., McLean, E. H., Steane, D. A., Potts, B. M., Vaillancourt, R. E., & Stock, W. D. (2015). Climate-adjusted provenancing: a strategy for climate-resilient ecological restoration. Frontiers in Ecology and Evolution, 3, 65.

R Development Core Team. (2020). R: A language and environment for statistical computing. R Foundation for Statistical Computing, Vienna, Austria. Retrieved from https://www.r-project.org/

Robson, T. M., Rasztovits, E., Aphalo, P. J., Alia, R., & Aranda, I. (2013). Flushing phenology and fitness of European beech (Fagus sylvatica L.) provenances from a trial in La Rioja, Spain, segregate according to their climate of origin. Agricultural and Forest Meteorology, 180, 76–85.

Runquist, R. B., Gorton, A. J., Yoder, J. B., Deacon, N. J., Grossman, J. J., Kothari, S., … Moeller, D. A. (2019). Context dependence of local adaptation to abiotic and biotic environments: a quantitative and qualitative synthesis. The American Naturalist, 707322.

Sinclair, F. H., Stone, G. N., Nicholls, J. A., Cavers, S., Gibbs, M., Butterill, P., … Schönrogge, K. (2015). Impacts of local adaptation of forest trees on associations with herbivorous insects: Implications for adaptive forest management. Evolutionary Applications, 8(10), 972–987.

Sun, M., Gross, K., & Schiestl, F. P. (2014). Floral adaptation to local pollinator guilds in a terrestrial orchid. Annals of Botany, 113(2), 289–300.

Vander Mijnsbrugge, K., Onkelinx, T., & De Cuyper, B. (2015). Variation in bud burst and flower opening responses of local versus non-local provenances of hawthorn (Crataegus monogyna Jacq.) in Belgium. Plant Systematics and Evolution, 301(4), 1171–1179.

Vázquez, D. P., Melián, C. J., Williams, N. M., Blüthgen, N., Krasnov, B. R., & Poulin, R. (2007). Species abundance and asymmetric interaction strength in ecological networks. Oikos, 116(7), 1120–1127.

Weißhuhn, K., Prati, D., Fischer, M., & Auge, H. (2012). Regional adaptation improves the performance of grassland plant communities. Basic and Applied Ecology, 13(6), 551–559.

Wilkinson, M., Eaton, E. L., & Morison, J. I. L. (2017). Variation in the date of budburst in Quercus robur and Q. petraea across a range of provenances grown in Southern England. European Journal of Forest Research, 136(1).

Williams, N. M., & Lonsdorf, E. V. (2018). Selecting cost-effective plant mixes to support pollinators. Biological Conservation, 217, 195–202.

Zelený, D., & Chytrý, M. (2019). Ecological specialization indices for species of the Czech flora. Preslia, 91(2), 93–116.

## References

Esser J, Fuhrmann M, Venne C (2020) Rote Liste und Gesamtartenliste der Wildbienen und Wespen – Hymenoptera – Aculeata – in Nordrhein-Westfalen, Landesamt für Natur, Umwelt und Verbraucherschutz Nordrhein-Westfalen, https://www.lanuv.nrw.de/natur/artenschutz/rote-liste/, last check 2020-02-24

Vázquez DP et al. (2007) Species abundance and asymmetric interaction strength in ecological networks. Oikos 116:1120–1127

Westrich P (2018) Die Wildbienen Deutschlands, Eugen Ulmer KG, Stuttgart, 821 pp.

